# Vacuole pH loss triggers ESCRT-dependent plasma membrane remodeling to prevent amino acid toxicity

**DOI:** 10.64898/2026.06.05.730386

**Authors:** Kevin C. Chui, Casey E. Hughes, Troy K. Coody, Adam L. Hughes

## Abstract

Loss of lysosomal or vacuolar acidity is a hallmark of aging, metabolic dysfunction, and cellular stress, yet how cells adapt to this condition remains poorly understood. In budding yeast, where the vacuole serves as a major reservoir for intracellular amino acids, impaired vacuolar acidification disrupts amino acid homeostasis. Here we performed a genome-wide screen in budding yeast to identify pathways required for survival during vacuole pH stress. We found that endocytic trafficking and ESCRT/MVB components become essential when vacuolar acidification is disrupted. Vacuole deacidification triggered ESCRT-dependent rerouting and degradation of plasma membrane amino acid transporters, thereby limiting nutrient influx. Blocking this response stabilized transporters at the cell surface and caused synthetic lethality under vacuole stress. This growth defect was suppressed by lowering amino acid availability or reducing transporter expression, whereas amino acid supplementation restored toxicity. Nitrogen starvation prevented transporter internalization, indicating that nutrient status gates this adaptive response. Together, these findings reveal a vacuole-plasma membrane communication pathway that protects cells from amino acid toxicity by matching nutrient influx to vacuolar function.

## INTRODUCTION

The vacuole in yeast and the lysosome in higher eukaryotes are acidic organelles that coordinate degradation, recycling, and nutrient storage (Abu-Remaileh et al., 2017; Feng et al., 2014; Jones et al., 1982; Kitamoto et al., 1988a; Kitamoto et al., 1988b; Klionsky et al., 1990; Klionsky et al., 1992; Lawrence and Zoncu, 2019; Nakamura et al., 1997; Weber et al., 2020; Wiemken and Dürr, 1974). Their acidic lumen enables proteolysis, autophagy (Nakamura et al., 1997), and ion and amino acid sequestration (Bianchi et al., 2019; Cai et al., 2023; Cools et al., 2020; Wang et al., 2014; Wiemken and Dürr, 1974), while their membranes integrate nutrient sensing with metabolic control (Demetriades et al., 2014; Rebsamen et al., 2015; Sancak et al., 2010; Wang et al., 2015). Through these functions, acidic compartments help maintain intracellular homeostasis and support adaptation to fluctuating metabolic conditions. In yeast, the vacuole is also a major storage depot for amino acids, phosphate, calcium, and metals, underscoring its broader role in buffering cytosolic nutrient composition during changing environmental conditions (Kitamoto et al., 1988a; Lander et al., 2016; Raguzzi et al., 1988; Wada et al., 1987; Wiemken and Dürr, 1974).

Disruption of vacuolar or lysosomal acidification therefore has widespread consequences for cellular physiology. In yeast, vacuole deacidification occurs naturally during aging and leads to mitochondrial fragmentation, iron dysregulation, and impaired respiratory function (Chen et al., 2020; Diab and Kane, 2013; Hughes and Gottschling, 2012; Hughes et al., 2020; Milgrom et al., 2007). More recently, vacuolar pH was shown to be dynamically regulated across the cell cycle, where programmed alkalinization and re-acidification help coordinate amino acid homeostasis with cell growth and division (Okreglak et al., 2023). In metazoans, lysosomal acidification defects contribute to diverse pathological conditions, including neurodegeneration, aging-associated decline, and cancer (Kim et al., 2023; Weber et al., 2020; Yambire et al., 2019). These observations highlight the importance of maintaining acidification for both metabolic homeostasis and cellular health.

Despite the broad physiological consequences of vacuole or lysosome dysfunction, cells can often tolerate acute perturbations in organelle acidity. Recent work from our laboratory showed that vacuole deacidification disrupts amino acid compartmentalization, causing cytosolic accumulation of specific amino acids that perturb iron homeostasis and mitochondrial function (Hughes et al., 2020). Work from other laboratories has likewise shown that the vacuole stores amino acids and metal ions in a highly regulated manner that responds to nutrient availability and cell cycle progression (Okreglak et al., 2023; Raguzzi et al., 1988; Russnak et al., 2001; Wada et al., 1987). These findings suggest that vacuole deacidification compromises the sequestration of metabolites that would otherwise be buffered away from the cytosol. Nevertheless, yeast cells remain viable following inhibition of the vacuolar H⁺-ATPase (V-ATPase), suggesting that compensatory mechanisms allow cells to adapt when vacuolar storage capacity is compromised. However, the adaptive responses that support survival during vacuole stress remain poorly understood.

Membrane trafficking pathways provide one potential mechanism by which cells adjust to metabolic stress. Endocytic trafficking dynamically regulates the abundance of plasma membrane nutrient transporters in response to environmental and metabolic cues (Edinger et al., 2009; García-Tardón et al., 2012; Kahlhofer et al., 2021; O’Donnell and Schmidt, 2019; Zbieralski and Wawrzycka, 2022). In yeast, the ubiquitin ligase Rsp5 and arrestin-related adaptor proteins promote ubiquitin-dependent internalization of permeases, while the ESCRT machinery mediates their sorting into multivesicular bodies for degradation in the vacuole (Babst, 2005; Helliwell et al., 2001; Henne et al., 2011; Lin et al., 2008; MacGurn et al., 2011; Nikko and Pelham, 2009; O’Donnell and Schmidt, 2019). Through this system, cells dynamically tune nutrient uptake in response to metabolic conditions. Despite extensive work on vacuolar nutrient storage and endocytic regulation of plasma membrane transporters, relatively little is known about how these systems are coordinated during vacuole stress. Notably, vacuolar and plasma membrane proton pumps cooperate to maintain pH homeostasis, raising the possibility that adaptive communication also links vacuolar function to transporter control at the cell surface (Martínez-Muñoz and Kane, 2008; Smardon and Kane, 2014; Velivela and Kane, 2018).

Here we performed a genome-wide screen to identify cellular processes required for survival during vacuole deacidification. Our results reveal that ESCRT-dependent plasma membrane trafficking becomes essential when vacuole acidity is disrupted. Loss of vacuole pH triggers widespread rerouting of amino acid transporters from the PM to the vacuole, and failure to remove these transporters leads to amino acid toxicity. Together, these findings uncover a vacuole–plasma membrane communication pathway that protects cells from nutrient overload during vacuole stress.

## RESULTS AND DISCUSSION

### Growth screen identifies ESCRT-dependent pathways required for survival during vacuole acidity loss

Vacuole deacidification disrupts multiple cellular processes, but how cells tolerate this condition remains unclear. To identify pathways required for adaptation to vacuole dysfunction, we screened the yeast haploid deletion collection (Giaever et al., 2002) for mutants that were hypersensitive to concanamycin A (concA), a vacuolar H⁺-ATPase (V-ATPase) inhibitor that deacidifies the vacuole (Fig. 1A). As previously described, concA slowed but did not completely block wild-type growth, enabling identification of genes that become conditionally essential during vacuole pH stress.

**Figure 1.**
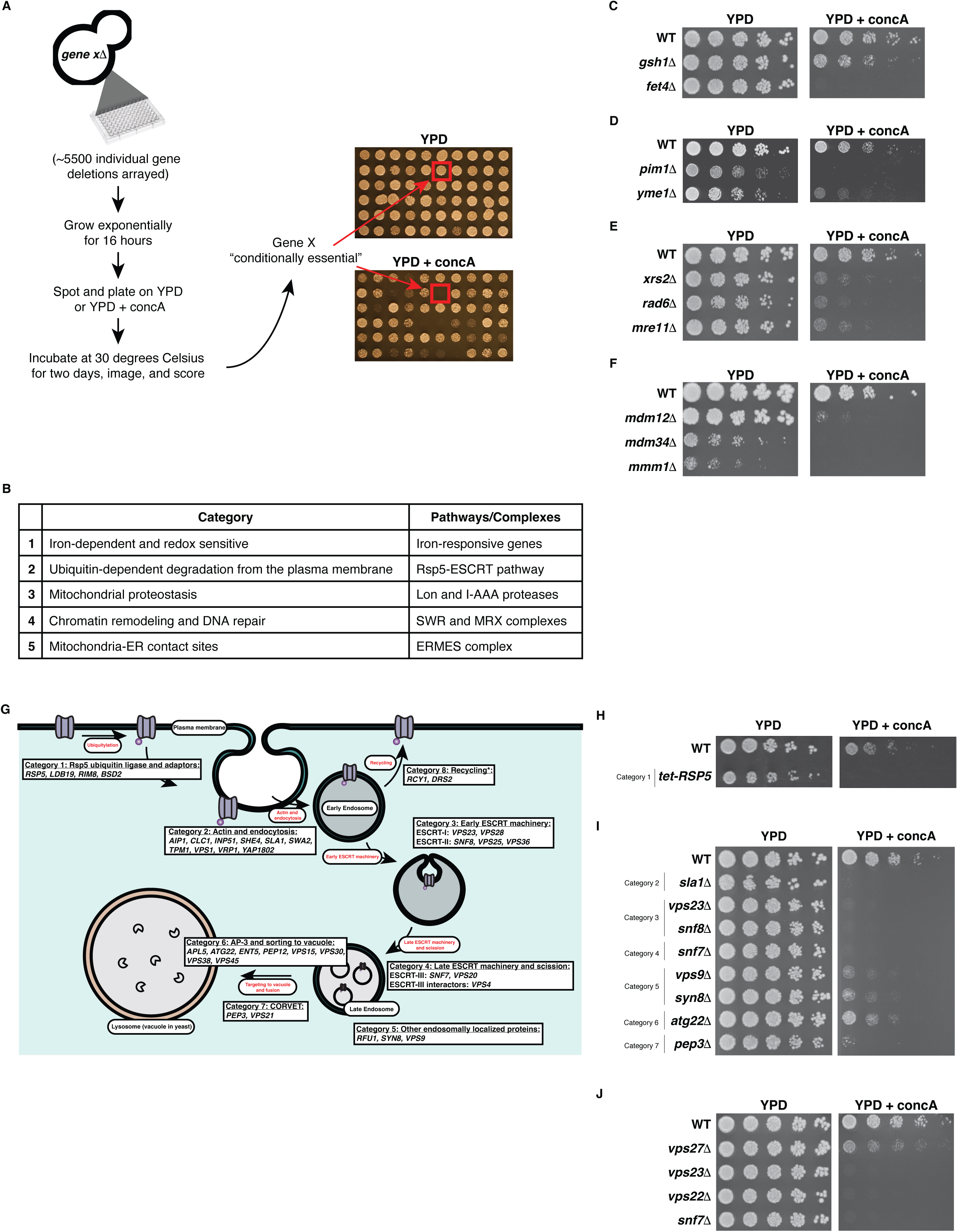
Genome-wide screen identifies pathways required for survival during vacuole deacidification. A. Schematic of the haploid deletion collection screen. Exponentially growing strains from the yeast haploid deletion collection were spotted onto YPD control or YPD + concA agar plates and grown for 2 days. Conditionally essential genes were identified based on reduced growth on YPD + concA relative to YPD control plates. B. Representative gene ontology analysis of conditionally essential genes identified in (A). A subset of functional categories were manually curated. C-F. Validation of representative screen hits related to categories in (B). Deletion mutants were reconstructed in BY4741 and analyzed by spot dilution growth assays on YPD or YPD + concA plates for 2 days. Representative genes from the following functional groups are shown: (C) iron/redox homeostasis, (D) mitochondrial proteases, (E) DNA repair, and (F) ERMES. G. Schematic summarizing genes across the endolysosomal trafficking pathway whose loss causes growth sensitivity in the presence of concA. All genes shown were identified in the genetic screen, with the exception of RSP5, which was tested directly based on related screen hits. H. Spot dilution growth assay using a tetracycline-repressible Rsp5 allele. I. Spot dilution growth assays retesting hits from multiple steps of the endolysosomal trafficking pathway described in (G). J. Validation of representative components from each ESCRT complex by spot dilution growth assay.

Two days after plating cells on solid rich medium with glucose (YPD) ± concA, colony growth was scored (see Materials and Methods) and sensitive mutants were identified and grouped into two categories (mild growth defect and severe growth defect; Fig. 1B; See Supplemental Table 1 for list of genes with alterations in growth with concA). Using gene ontology (GO)-term analysis, several pathways were enriched among concA-sensitive mutants—examples include genes involved in iron and redox homeostasis (Hughes et al., 2020; Stearman et al., 1996), mitochondrial proteostasis (Van Dyck and Langer, 1999), chromatin remodeling and DNA repair (Jentsch et al., 1987; Prakash, 1974), and mitochondria–ER contact sites (English et al., 2020; Kornmann et al., 2011; Schuler et al., 2021). Representative mutants from these categories—including iron/redox (*gsh1Δ*, *fet4Δ*), mitochondrial proteases (*pim1Δ*, *yme1Δ*), DNA repair (*xrs2Δ*, *rad6Δ*, *mre11Δ*), and ERMES components (*mdm12Δ*, *mdm34Δ*, *mmm1Δ*)—displayed clear sensitivity to vacuole pH disruption (Fig. 1C–F).

Among the most consistently recovered hits were genes involved in endolysosomal trafficking and the Endosomal Sorting Complex Required for Transport (ESCRT)/multivesicular body (MVB) pathway, which, as described above, regulates the abundance of nutrient transporters at the cell surface (Babst, 2005; Henne et al., 2011). To further examine this class, we expanded our analysis across multiple steps of plasma membrane remodeling and endosomal trafficking. Sensitive mutants were identified in pathways mediating cargo recognition and ubiquitin-dependent sorting (Gajewska et al., 2003), actin-dependent endocytosis (Holtzman et al., 1993; Na et al., 1995), early ESCRT complexes (Babst et al., 2000; Rothman et al., 1989), late ESCRT components responsible for membrane scission (Babst et al., 1998), as well as additional factors involved in endosomal sorting (Burd et al., 1996; Robinson et al., 1988), vacuolar trafficking (Cowles et al., 1997), and endosome–vacuole fusion and recycling (Robinson et al., 1988; Rothman et al., 1989) (Fig. 1G–J). Representative mutants from each ESCRT complex—including *vps27Δ* (ESCRT-0), *vps23Δ* (ESCRT-I), *vps22Δ* (ESCRT-II), and *snf7Δ* (ESCRT-III)—showed strong growth defects in the presence of concA. Altogether, these results indicate that multiple adaptive pathways contribute to survival when vacuole acidity is perturbed. Notably, the strong requirement for endocytic and ESCRT/MVB components suggested that membrane trafficking plays a central role in the cellular response to vacuole stress.

### Vacuole acidity loss triggers ESCRT-dependent rerouting of plasma-membrane transporters

Because our screen identified ESCRT/MVB components as conditionally essential during vacuole acidification stress, we next asked whether vacuole deacidification triggers adaptive remodeling of the plasma membrane. This possibility is supported by earlier studies showing that V-ATPase dysfunction perturbs plasma membrane homeostasis, including crosstalk with the proton pump Pma1, and that cells lacking V-ATPase activity require endocytosis and Rsp5-dependent trafficking for survival (Munn and Riezman, 1994; Smardon and Kane, 2014). Because the vacuole is a major intracellular reservoir for amino acids, we reasoned that loss of vacuole acidity might compromise amino acid buffering, creating a need to limit further influx at the cell surface. We therefore hypothesized that cells adapt to vacuole stress by limiting the accumulation of plasma membrane amino acid transporters at the cell surface.

To test this idea, we examined the localization of amino acid permeases following inhibition of vacuole acidification with concA. Using yeast strains expressing GFP-tagged transporters, we first used immunoblotting to determine which amino acid transporters were detectably expressed under these conditions (Supplemental Fig.1). Several transporters were readily detected, and we chose to focus initially on Bap2, a branched-chain amino acid permease as a representative cargo (Grauslund et al., 1995). Under normal exponential growth conditions, Bap2–GFP localized primarily to the plasma membrane with some signal in the vacuole lumen (Fig. 2A). After 3 hours of concA treatment, Bap2 was internalized and accumulated within the vacuole lumen, consistent with delivery for degradation. Immunoblot analysis confirmed this remodeling, showing a reduction in full-length Bap2–GFP and accumulation of free GFP, which reflects vacuolar degradation of the transporter (Fig. 2B). Notably, transporter degradation still occurred despite inhibition of vacuole acidification, consistent with previous studies showing that concA treatment does not fully block vacuolar proteolysis in this strain background (Schuler et al., 2021).

**Figure 2.**
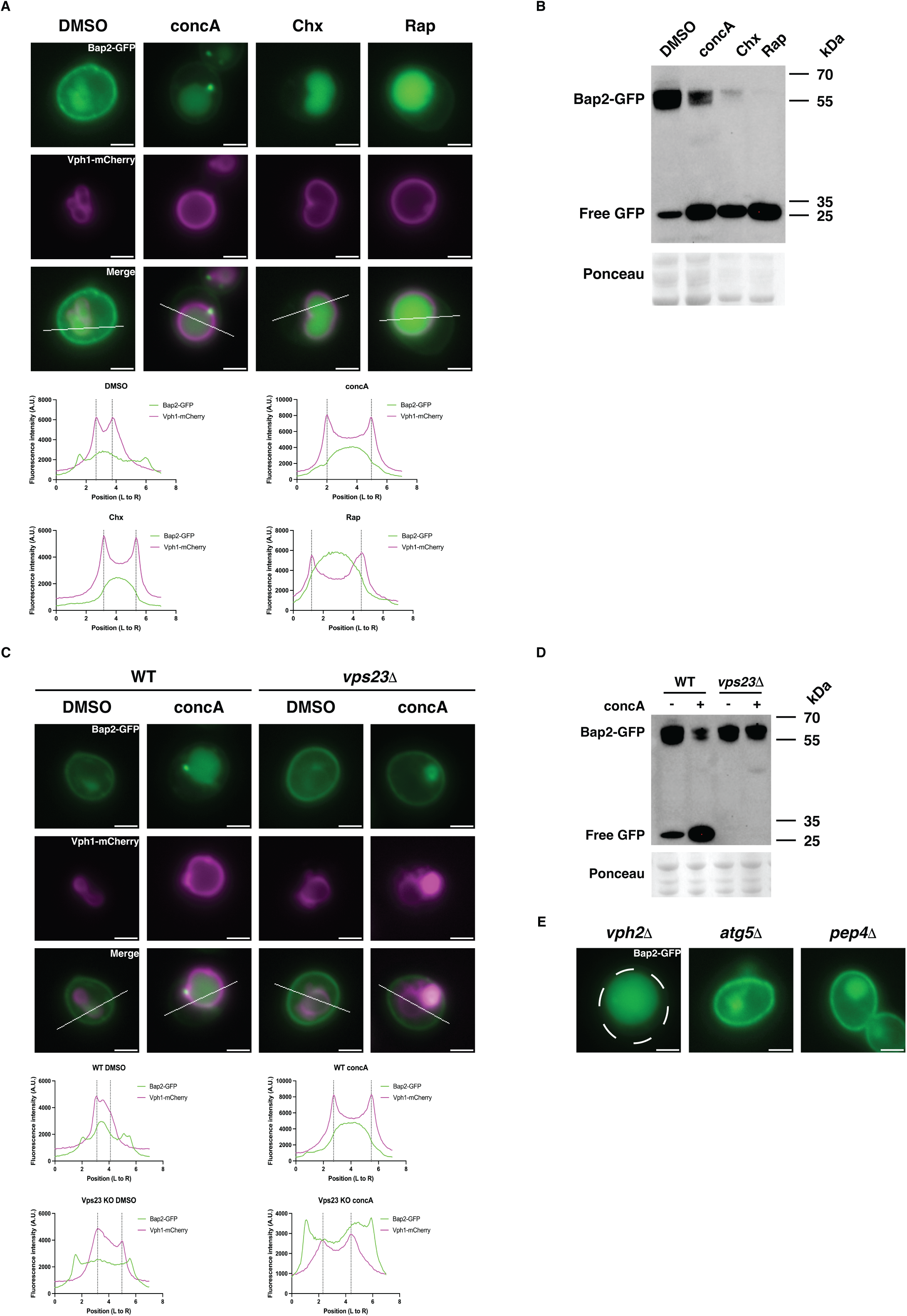
Vacuole deacidification triggers ESCRT-dependent rerouting of amino acid transporters to the vacuole. A. Representative images of Bap2–GFP following treatment with DMSO, concA, cycloheximide (CHX), or rapamycin (Rap), with accompanying line scans. Cells were grown to exponential phase, treated for 3 h, and imaged. Vacuole membranes were labeled with Vph1–mCherry. Scale bars: 2 µm. See Methods for microscopy reproducibility and validation details. B. Western blot analysis of Bap2–GFP following treatment with DMSO, concA, CHX, or Rap for 3 h. Cells were harvested from exponentially growing cultures. Primary antibody: α-GFP. C. Representative images of Bap2–GFP in WT or *vps23Δ* cells following 3 h treatment with DMSO or concA. Scale bars: 2 µm. D. Western blot analysis of Bap2–GFP following 3 h treatment with DMSO or concA in WT, *vps23Δ*, and additional ESCRT mutant strains. Primary antibody: α-GFP. E. Representative images of Bap2–GFP localization in mutants defective in V-ATPase assembly (*vph2Δ*), autophagy (*atg5Δ*), or vacuolar proteolysis (*pep4Δ*). Cells were grown to exponential phase and imaged. Scale bars: 2 µm.

Drugs known to induce endocytic turnover of amino acid permeases, including the translation inhibitor cycloheximide and the mTOR inhibitor rapamycin, produced similar transporter rerouting and degradation patterns on the same timescale as concA treatment (Fig. 2A-B) (Lin et al., 2008; MacGurn et al., 2011; Omura and Kodama, 2004). Importantly, concA-induced internalization also required ESCRT function. Transporter routing to the vacuole was blocked in individual mutants lacking proteins from the various ESCRT complexes (Fig. 2C-D; Supplemental Fig. 1B). In contrast, deletion of the vacuolar protease Pep4 (Ammerer et al., 1986; Jones et al., 1982), which acts downstream of the ESCRT/MVB pathway, did not impair transporter delivery to the vacuole, indicating that ESCRT-dependent trafficking rather than vacuolar proteolysis is required for entry to the vacuole lumen (Supplemental Fig. 1B).

To determine whether transporter internalization specifically reflects defects in vacuole acidification, we examined mutants affecting vacuolar proteolysis or autophagy, which disrupt vacuolar protein degradation but do not alter pH (Feng et al., 2014; Jones et al., 1982). Loss of Pep4 (Ammerer et al., 1986; Jones et al., 1982) or the core autophagy component Atg5 (Feng et al., 2014) did not trigger Bap2 vacuole delivery, whereas deletion of Vph2—which disrupts V-ATPase assembly and vacuolar acidification (Preston et al., 1992) —was sufficient to trigger constitutive Bap2 turnover (Fig. 2E). These results indicate that defective vacuole acidification triggers ESCRT-dependent degradation of plasma membrane transporters, indicating that transporter turnover is triggered by vacuolar acidification defects rather than by impaired vacuolar proteolysis itself.

Finally, we asked whether this response extends beyond Bap2 to other transporters expressed in nutrient-rich medium. Several additional amino acid permeases, including Dip5, Lyp1, and Ptr2, were similarly rerouted from the PM to the vacuole and degraded following vacuole pH disruption (Supplemental Fig. 1C-D). Together, these findings demonstrate that vacuole deacidification triggers broad ESCRT-dependent rerouting of amino acid transporters to the vacuole, suggesting an adaptive response that limits amino acid influx when vacuolar function is compromised.

### Synthetic lethality between vacuole deacidification and loss of ESCRT function is driven by amino acid load

Because vacuole deacidification triggered ESCRT-dependent rerouting of plasma membrane transporters to the vacuole (Fig. 2), we next tested whether amino acid influx contributes to the synthetic lethality observed between concA treatment and loss of ESCRT function. If transporter rerouting protects cells from amino acid overload, reducing extracellular amino acids should alleviate the growth defect, whereas restoring amino acid supply should reinstate toxicity. To test this idea, we compared growth of wild-type and ESCRT mutants in rich medium (YPD) and synthetic defined medium (SD), the latter of which contains lower levels of amino acids (Schuler et al., 2021). We found that ESCRT mutants—including representatives from each complex (*vps27Δ*, *vps23Δ*, *vps22Δ*, *snf7Δ*)—grew normally on SD ± concA (Fig. 3A). Thus, lowering amino acid availability suppresses the synthetic lethality between vacuole pH loss and ESCRT disruption. We also found that iron supplementation rescued growth of ESCRT mutants in YPD + concA (Fig. 3A). This observation is consistent with our previous work showing that vacuole deacidification disrupts amino acid compartmentalization and leads to secondary iron limitation and mitochondrial dysfunction (Hughes et al., 2020). However, because lowering amino acid availability alone restored growth, these results suggest that excess amino acid influx is the primary driver of toxicity when ESCRT-dependent trafficking is impaired in V-ATPase-deficient cells.

**Figure 3.**
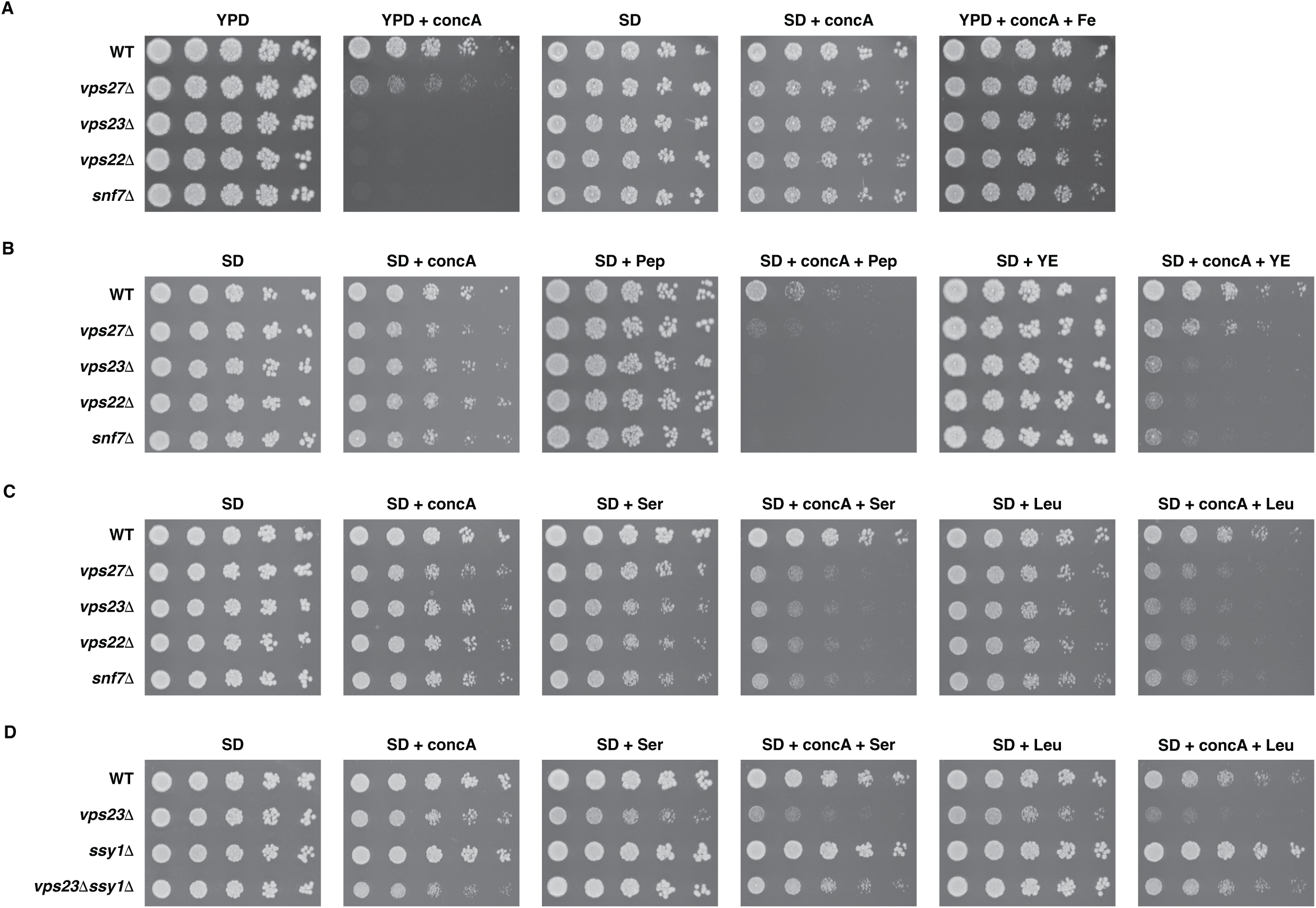
Amino acid availability drives synthetic lethality between vacuole deacidification and ESCRT disruption. A. Spot dilution growth assays of WT and ESCRT mutant strains grown on rich (YPD) or low amino acid synthetic (SD) medium containing DMSO, concA, or iron supplementation. B. Spot dilution growth assays of WT and ESCRT mutant strains grown on SD medium supplemented with concA and/or peptone (Pep) or yeast extract (YE). C. Spot dilution growth assays of WT and ESCRT mutant strains grown on SD medium containing DMSO or concA and supplemented with serine (35 mM) or leucine (75 mM). D. Spot dilution growth assays of WT, *vps23Δ*, *ssy1Δ*, and *vps23Δ ssy1Δ* strains grown on SD medium containing DMSO or concA and supplemented with serine (35 mM) or leucine (75 mM).

We next asked whether restoring amino acid sources would reestablish synthetic lethality. Addition of bulk amino acid sources such as peptone or yeast extract restored concA sensitivity in ESCRT mutants grown in SD medium (Fig. 3B). Supplementation of SD with higher levels of individual amino acids—including serine, leucine, arginine, or cysteine—produced a similar effect (Fig. 3C; Supplemental Fig. 2A). These results suggest that amino acid availability directly drives the growth defect observed when ESCRT-dependent trafficking is compromised, and that multiple amino acids contribute to this growth impairment.

If this toxicity results from a failure of ESCRT mutants to reroute amino acid transporters away from the plasma membrane, then reducing transporter expression should suppress the phenotype. However, deletion of a number of individual permeases in *vps23Δ* cells did not restore growth on YPD + concA (Supplemental Fig. 2B), likely because of redundancy among the more than twenty amino acid transporters at the plasma membrane and because multiple amino acids are toxic under these conditions (Fig. 3C; Supplemental Fig. 2A) (Bianchi et al., 2019). We therefore turned to the Ssy1–Ptr3–Ssy5 (SPS) amino acid sensing pathway, which coordinately regulates expression of multiple transporters in response to extracellular amino acids (Forsberg et al., 2001; Klasson et al., 1999). *SSY1* encodes the plasma membrane sensor for this pathway and is required for expression of several SPS-regulated transporters (Forsberg et al., 2001; Klasson et al., 1999). Deletion of Ssy1 alone did not rescue growth of ESCRT mutants on YPD + concA (Supplemental Fig. 2B), likely because multiple amino acids contribute to toxicity in this condition and not all relevant transporters are controlled by the SPS system. In contrast, loss of Ssy1 restored growth of *vps23Δ* cells on SD + concA medium supplemented with individual amino acids such as serine, leucine, or cysteine, although this suppression was not observed for arginine (Fig. 3D; Supplemental Fig. 2C). This lack of rescue with arginine is consistent with the substrate range of SPS-regulated permeases, many of which do not transport basic amino acids (Forsberg et al., 2001; Klasson et al., 1999). Together, these results indicate that the synthetic lethality between vacuole pH loss and ESCRT disruption is driven by excessive amino acid influx, supporting a model in which ESCRT-dependent transporter rerouting protects cells when vacuole amino acid compartmentalization is compromised.

### Nitrogen and amino acid availability control transporter rerouting during vacuole pH loss

Finally, we sought to identify the signal that triggers ESCRT-dependent rerouting of plasma membrane transporters when vacuole acidification is impaired. This question was important because previous studies demonstrated that changes in vacuolar pH can drive signaling between the plasma membrane proton pump Pma1 and the V-ATPase, indicating vacuole pH changes can influence plasma membrane physiology (Velivela and Kane, 2018). Because the transporters affected in our experiments primarily mediate amino acid uptake, we hypothesized that amino acid availability or nitrogen status might contribute to the signal that promotes transporter internalization (Chen and Kaiser, 2002). To test this possibility, we examined whether transporter internalization during concanamycin A (concA) treatment depended on extracellular amino acid levels. Notably, the synthetic lethality between concA treatment and loss of ESCRT function was suppressed when cells were grown in low–amino acid synthetic defined (SD) medium (Fig. 3A), suggesting that amino acid availability influences the requirement for ESCRT function under these conditions. However, Bap2–GFP was still robustly internalized following concA treatment in SD medium (Fig. 4A-B), indicating that the signal triggering transporter rerouting persists even when extracellular amino acids are reduced.

**Figure 4.**
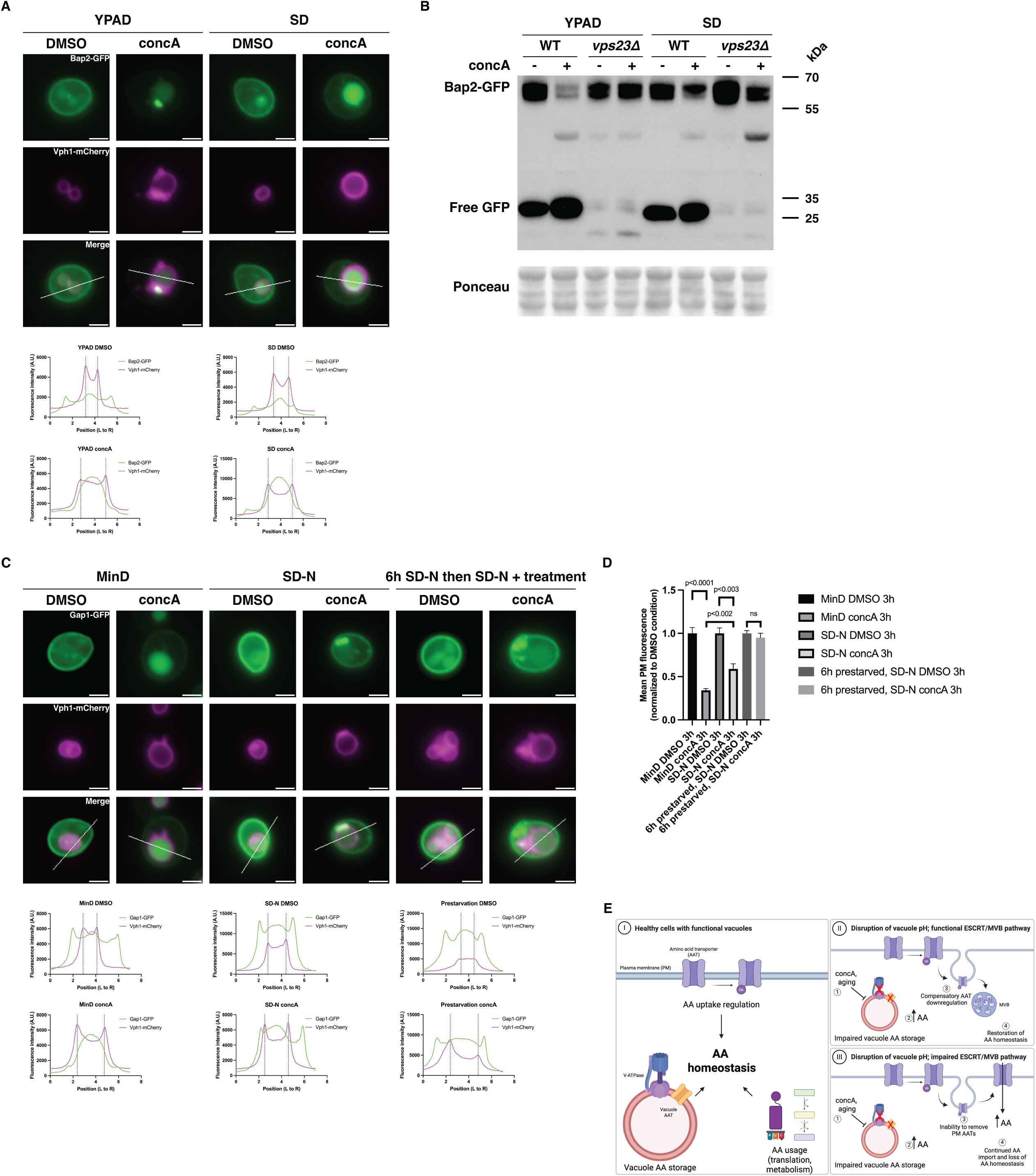
Nitrogen and amino acid availability regulate transporter rerouting during vacuole stress. A. Representative images of Bap2–GFP following 3 h treatment with DMSO or concA in YPD or SD medium. Scale bars: 2 µm. B. Western blot analysis of Bap2–GFP following 3 h treatment with DMSO or concA in YPD or SD medium in WT and *vps23Δ* strains. Primary antibody: α-GFP. C. Microscopy of Gap1–GFP following 3 h treatment with DMSO or concA under minimal medium (MinD), nitrogen starvation (SD–N), or nitrogen prestarvation conditions (6 h SD–N followed by SD–N). Scale bars: 2 µm. D. Quantification of plasma membrane GFP fluorescence intensity for cells shown in (C). Plasma membrane GFP intensities were measured from ≥100 cells per condition and averaged for each replicate prior to normalization to the corresponding DMSO control within each media condition. Data represent n = 3 biological replicates. E. Model for adaptive plasma membrane remodeling during vacuole stress. (I) Amino acid homeostasis is maintained through regulated uptake, intracellular compartmentalization, and utilization. (II) Loss of vacuolar amino acid compartmentalization triggers compensatory downregulation of plasma membrane amino acid transporters. (III) Simultaneous impairment of vacuolar amino acid compartmentalization and transporter downregulation disrupts amino acid homeostasis and promotes amino acid toxicity.

Iron supplementation also rescued growth of ESCRT mutants during vacuole stress (Fig. 3A), raising the possibility that iron limitation might regulate this response. However, addition of iron did not block concA-induced Bap2 rerouting, and iron chelation alone with bathophenanthroline disulphonate (BPS) was insufficient to trigger changes in Bap2-GFP localization (Supplemental Fig. 3A-B). Similarly, treatment with the antioxidant glutathione (GSH) did not prevent transporter rerouting (Supplemental Fig. 3C-D). These observations indicate that neither iron availability nor oxidative stress that arises downstream of iron perturbation drives the changes in transporter location, pointing again to amino acid or nitrogen cues as potential upstream signals.

To test this possibility more directly, we examined transporter dynamics under defined nitrogen conditions. In synthetic minimal medium (MinD), ammonium sulfate provides nitrogen in the absence of extracellular amino acids. Under these conditions, cells induce expression of the general amino acid permease Gap1 while repressing specific permeases such as Bap2 (Magasanik and Kaiser, 2002). Because Bap2 expression is limited in MinD, we used Gap1–GFP as a reporter to monitor plasma membrane remodeling under nitrogen-controlled conditions.

In MinD, Gap1–GFP localized primarily to the plasma membrane with modest vacuolar signal reflecting basal turnover (Fig. 4C-D). ConcA treatment triggered strong rerouting and degradation of Gap1–GFP, demonstrating that vacuole deacidification is sufficient to induce transporter rerouting even when extracellular amino acids are absent (Fig. 4C-D). We next tested whether nitrogen starvation alters this response. When MinD-grown cells were transferred to nitrogen-free medium (SD–N), concA-induced Gap1 rerouting was substantially reduced, with transporter signal remaining at the plasma membrane in many cells (Fig. 4C-D). Quantification showed that plasma membrane fluorescence decreased to ∼25% of control levels in MinD but only to ∼50% in SD–N (Fig. 4D). We reasoned that residual internalization after the nitrogen shift might reflect intracellular amino acid pools that persist transiently (Wiemken and Dürr, 1974). To deplete these pools, cells were pre-starved for six hours in SD–N before concA treatment. Under these conditions, concA treatment failed to trigger detectable Gap1 rerouting and degradation (Fig. 4C–D). Together, these experiments suggest that vacuole pH–induced transporter rerouting to the vacuole requires amino acid and nitrogen availability. The response is not driven by iron limitation or oxidative stress and is suppressed when intracellular amino acid pools are depleted. These findings support a model in which altered amino acid or nitrogen status in V-ATPase-impaired cells promotes ESCRT-dependent plasma membrane remodeling, thereby limiting amino acid influx when nutrient toxicity is likely to arise.

### Model and implications

The vacuole (or lysosome in mammals), once viewed primarily as a degradative compartment, is increasingly recognized as a central hub for nutrient sensing and metabolic regulation (Amick et al., 2020; Cools et al., 2020; Hughes and Gottschling, 2012; Hughes et al., 2020; Lawrence and Zoncu, 2019; Okreglak et al., 2023; Rebsamen et al., 2015). Previous work from Kane and colleagues demonstrated that vacuole acidification can influence plasma membrane composition by regulating turnover of the proton pump Pma1 through cytosolic pH–dependent mechanisms (Smardon and Kane, 2014; Velivela and Kane, 2018). Our findings extend this concept by identifying a parallel remodeling pathway that targets amino acid transporters. When vacuole acidity is lost, cells activate an ESCRT-dependent remodeling program that reroutes plasma membrane amino acid transporters to the vacuole, thereby limiting nutrient influx. This response protects cells from amino acid toxicity and supports survival during organelle stress. Our screen also identified additional pathways—including mitochondrial proteostasis, chromatin remodeling, and DNA repair—that become important when vacuole acidity is disrupted, suggesting that vacuole deacidification triggers broader cellular adaptation. These processes likely help cells cope with metabolic and oxidative stresses associated with disrupted amino acid and iron homeostasis (Hughes et al., 2020; Yambire et al., 2019).

Based on our results, we propose a model (Fig. 4E) in which vacuole pH loss leads to miscompartmentalization of amino acids and secondary metabolic stress as previously described (Hughes et al., 2020; Schuler et al., 2021). Under nutrient-replete conditions, this imbalance triggers ESCRT-dependent endocytosis of plasma membrane amino acid transporters via the Rsp5–arrestin network, reducing amino acid influx and preventing amino acid toxicity. In contrast, nitrogen starvation suppresses this response, indicating that amino acid and nitrogen availability gate vacuole-to-plasma membrane signaling. In this way, cells coordinate plasma membrane transporter activity with vacuolar amino acid compartmentalization to maintain intracellular amino acid homeostasis. Although the exact molecular signals linking vacuole deacidification to PM remodeling remain to be defined, our results suggest that intracellular amino acid or nitrogen status acts as a driver in V-ATPase-deficient cells. Together, these findings reveal a principle of cellular adaptation in which organelle dysfunction triggers coordinated remodeling of the plasma membrane to rebalance nutrient flux and maintain metabolic homeostasis. Our results further underscore that intracellular amino acid levels must be tightly regulated to prevent toxicity, extending prior work from our laboratory and others (Hughes et al., 2020; Ohashi et al., 2017; Risinger et al., 2006; Ruiz et al., 2020; Schuler et al., 2021), and reveal coordinated communication between the vacuole and plasma membrane that safeguards cells from nutrient overload.

## MATERIALS AND METHODS

### Yeast strain construction

All strains (Supplemental Table 2) were derived from *Saccharomyces cerevisiae* S288C (Brachmann et al., 1998). Gene deletions were performed using one-step PCR-mediated gene replacement with the pRS vector series (oligos listed in Supplemental Table 3; plasmids in Supplemental Table 4). Strains with GFP-tagged proteins were constructed with either pKT127 (KanMX selection) or pKT209 (*URA3* selection). Strains with mCherry-tagged Vph1 were constructed with pFA6a-mCherry-HYG (HygMX selection). All strains were endogenously tagged at their C-termini. PCR products were integrated at the endogenous locus by replacing the native stop codon with the fluorescent tag followed by a new stop codon, thereby maintaining expression from the native promoter. Correct integrations and gene deletions for all strains were confirmed by colony PCR across the chromosomal insertion site and, where appropriate, microscopy visualizing the correct subcellular localization of the fusion protein.

### Cell culture

Unless otherwise noted, yeast were grown at 30°C in rich, high amino acid medium supplemented with glucose (YPD), which contained 1% yeast extract, 2% peptone, and 0.005% adenine. Low amino acid synthetic medium (S) contained 0.67% yeast nitrogen base without amino acids, 0.074 g/L each adenine, alanine, arginine, asparagine, aspartic acid, cysteine, glutamic acid, glutamine, glycine, histidine, myo-inositol, isoleucine, lysine, methionine, phenylalanine, proline, serine, threonine, tryptophan, tyrosine, uracil, valine, 0.369 g/L leucine, and 0.007 g/L para-aminobenzoic acid. Experiments performed with minimal medium (Min) contained 0.67% yeast nitrogen base without amino acids. Nitrogen starvation medium (S-N) contained 0.17% yeast nitrogen base without amino acids or ammonium sulfate. Glucose (D) was added at 2% for all media. For addback experiments, peptone and yeast extract were added to synthetic medium at final concentrations of 2% and 1%, respectively.

For each experiment, yeast strains were recovered from freezer stock on agar plates at 30°C overnight and used to inoculate liquid cultures that were saturated at 30°C overnight. For YPD conditions, 0.5 µL saturated culture was inoculated in 50 mL fresh media and grown exponentially overnight to a final density of 1-4 × 10^6^ cells/mL. For SD and MinD conditions, 1 µL and 15 µL saturated culture was used, respectively. Final concentrations for drug treatments were concanamycin A (concA; 500 nM), ferrous ammonium sulfate (denoted as Fe or iron; 2 mM), bathophenanthroline disulphonate (BPS; 250 µM), cycloheximide (Chx; 50 µg/mL), rapamycin (Rap; 200 ng/mL), and glutathione (GSH; 2 mM). Unless otherwise noted, treatments were performed for 3 h.

### Haploid deletion collection screen

The Yeast Haploid Deletion Collection (Dharmacon) was pinned onto YPD agar plates and grown for 24 h. Colonies were transferred to 96-well plates containing YPD and grown to saturation for 2 days. To generate log-phase cultures, 1 µL of each saturated culture was inoculated into 200 µL YPD and grown for 4 h. Aliquots of log-phase cultures (2 µL) were then spotted onto YPD plates in the presence or absence of concanamycin A (concA; 500 nM), and growth was scored after 2 days. Sensitivity was scored by comparing growth on YPD + concA plates relative to YPD control plates using a three-category scale: not sensitive, mild growth defect, and severe growth defect. Gene ontology enrichment analysis was performed on the set of concA-sensitive mutants using g:Profiler (https://biit.cs.ut.ee/gprofiler/gost) as an exploratory tool to identify candidate pathways for follow-up analysis.

### Spot dilution growth assays

This was performed as previously described (Hughes et al., 2020). Yeast strains were recovered from freezer stocks on YPD agar plates for 24 hours. They were then inoculated in 4 mL of YPD liquid media in culturing tubes and grown to saturation for 24 hours. The following day, 100 µL of saturating culture was added to 8 mL of fresh YPD. Yeast were grown to exponential phase for three hours and then diluted with water to normalize cell densities across strains. Five-fold serial dilutions were made in water, and aliquots (3 µL) of each dilution were spotted onto agar plates, resulting in approximately 5000, 1000, 200, 40, and 8 cells per spot.

### Microscopy and image analysis

Unless otherwise noted, overnight log-phase cell cultures were grown in the presence of dimethyl sulfoxide (DMSO) or the indicated treatment for three hours. After incubation, cells were harvested by centrifugation, resuspended in imaging buffer (5 % Glucose, 10 mM HEPES pH 7.6) and plated onto a slide to allow for the formation of a monolayer.

Optical z-sections of live yeast cells were acquired with a Zeiss Axiocam 506 monochromatic camera and 63 x oil immersion objectives (Carl Zeiss, Plan Apochromat, NA 1.4). To obtain super-resolution images a ZEISS LSM800 confocal microscope equipped with an Airyscan detector and 63 x oil immersion objective (Carl Zeiss, Plan Apochromat, NA 1.4) was used. Images were processed and analyzed using a combination of Zen software (Carl Zeiss) and Fiji. All representative images are from a single focal plane.

### Microscopy reproducibility and validation

All microscopy experiments were independently repeated at least three times with similar results. Representative images are shown from analyses of>50 cells, and the displayed localization patterns were representative of the majority of cells observed under each condition. Where indicated, microscopy-based localization phenotypes were validated using orthogonal GFP cleavage and immunoblot assays from population-level samples.

### Quantification of mean plasma membrane GFP intensity

Raw images were processed in Fiji. Regions of interest (ROIs) were drawn along the plasma membrane using the segmented line tool, and mean GFP fluorescence intensity was measured for each ROI. Measurements were collected from all cells across all images. For each replicate and condition, mean plasma membrane fluorescence intensity was calculated and normalized to the corresponding DMSO control within the same media condition. Normalized values were plotted as bar graphs. Statistical significance between DMSO and concA-treated samples within each media condition was determined using two-tailed t-tests.

### Western blotting

Yeast were grown to exponential phase overnight and treated as indicated for each experiment. A total of 2 × 10⁷ cells were harvested, washed in 1000 µL water, and resuspended in 100 µL 100 mM NaOH for 5 min at room temperature. Cells were then centrifuged at 4°C for 10 min at 20,000 × g. The cells were resuspended in SDS lysis buffer (10 mM Tris-HCl pH 6.8, 100 mM NaCl, 1% SDS, 1 mM EGTA, 1 mM EDTA, 1x cOmplete protease inhibitors) and SDS-PAGE loading buffer (30 mM Tris-HCl pH 6.8, 3% SDS, 5% glycerol, 0.004% bromophenol blue, 2.5% β-mercaptoethanol). Samples were incubated at 50°C for 15 min, placed on ice for 30 s, and centrifuged for 3 min at 20,000 × g. Supernatants were resolved on Bolt 4–12% Bis-Tris Plus gels with MES/SDS buffer and transferred to nitrocellulose membranes with the Invitrogen Power Blotter Station. Membranes were stained with Ponceau S and imaged. Membranes were then blocked and probed in blocking buffer (TBS; 50 mM Tris-HCl pH 7.4, 150 mM NaCl, 0.05% Tween-20, 5% non-fat dry milk) using mouse α-GFP primary antibodies and HRP-conjugated secondary antibodies. Blots were developed using Dura Supersignal Enhanced Chemiluminescence Solution and imaged on a ChemiDoc MP system.

## ONLINE SUPPLEMENTAL MATERIAL

Supplemental Fig.1 shows microscopy and GFP clipping assays for additional amino acid transporters examined in this study. Supplemental Fig. 2 shows spot assays of ESCRT mutants grown on arginine- or cysteine-supplemented media, as well as single amino acid transporter deletions in the model ESCRT mutant *vps23Δ* grown on YPD and YPD + concA. Supplemental Fig. 3 shows Bap2–GFP microscopy assays following treatment with iron supplementation, the iron chelator BPS, or glutathione.

Supplemental Table 1 lists gene hits identified in the concA screen. Supplemental Table 2 lists the strains used in this study. Supplemental Table 3 lists the oligos used in this study. Supplemental Table 4 lists the reagents and plasmids used in this study.

## DATA AVAILABILITY

All reagents used in this study are available upon request. All other data reported in this paper will be shared by the lead contact upon request. This paper does not report original code. Any additional information required to reanalyze the data reported in this paper is available from the lead contact upon request.

## ACKNOWLEDGEMENTS

We thank past and present members of the A.L. Hughes group for discussion and manuscript comments. Research was supported by National Institutes of Health grants GM119694 and AG061376 to A.L.H, NIH T32 DK091317 to T.K.C., NIH T32 DK007115 to C.E.H., and NIH F31 AG082494 to K.C.C. The content is solely the responsibility of the authors and does not necessarily represent the official views of the NIH.

## AUTHOR CONTRIBUTIONS

Conceptualization, K.C.C., T.K.C., C.E.H., and A.L.H.; methodology, K.C.C., T.K.C., C.E.H., and A.L.H.; formal analysis, K.C.C., T.K.C., and C.E.H.; investigation, K.C.C., T.K.C., and C.E.H; writing – original draft, K.C.C.; writing – review and editing, K.C.C. and A.L.H.; visualization, K.C.C.; supervision, A.L.H.; funding acquisition, K.C.C., T.K.C., C.E.H., and A.L.H.

## CONTACT FOR REAGENT AND RESOURCE SHARING

Further information and requests for resources and reagents should be directed to and will be fulfilled by the Lead Contact, Adam Hughes. All unique/stable reagents generated in this study are available from the Lead Contact without restrictions.

## SUPPLEMENTAL FIGURE LEGENDS

**Supplemental Figure 1.**
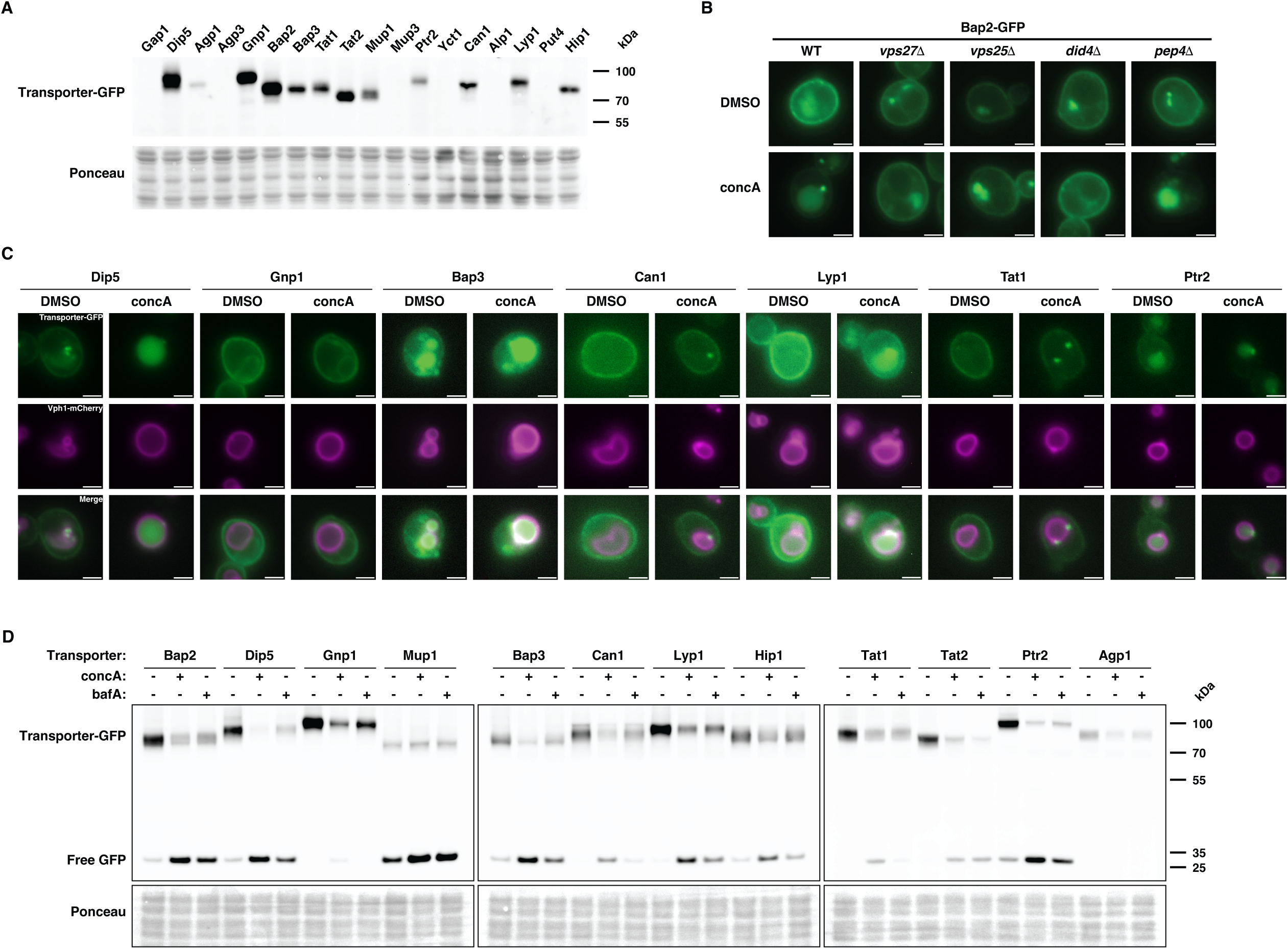
Vacuole deacidification triggers broad rerouting of plasma membrane amino acid transporters to the vacuole (related to Figure 2) (A) Western blot analysis of exponentially growing strains expressing GFP-tagged plasma membrane amino acid transporters (AATs) in YPD medium. Cells were harvested from exponentially growing cultures. Primary antibody: α-GFP. (B) Microscopy of Bap2–GFP localization following 3 h treatment with DMSO or concA in WT, ESCRT mutant, or *pep4Δ* strains. Scale bars: 2 µm. (C) Microscopy showing localization of additional GFP-tagged plasma membrane amino acid transporters following 3 h treatment with DMSO or concA. Scale bars: 2 µm. (D) Western blot analysis of GFP-tagged plasma membrane amino acid transporters following 3 h treatment with DMSO, concA, or bafilomycin A1 (bafA). Cells were harvested from exponentially growing cultures. Primary antibody: α-GFP.

**Supplemental Figure 2.**
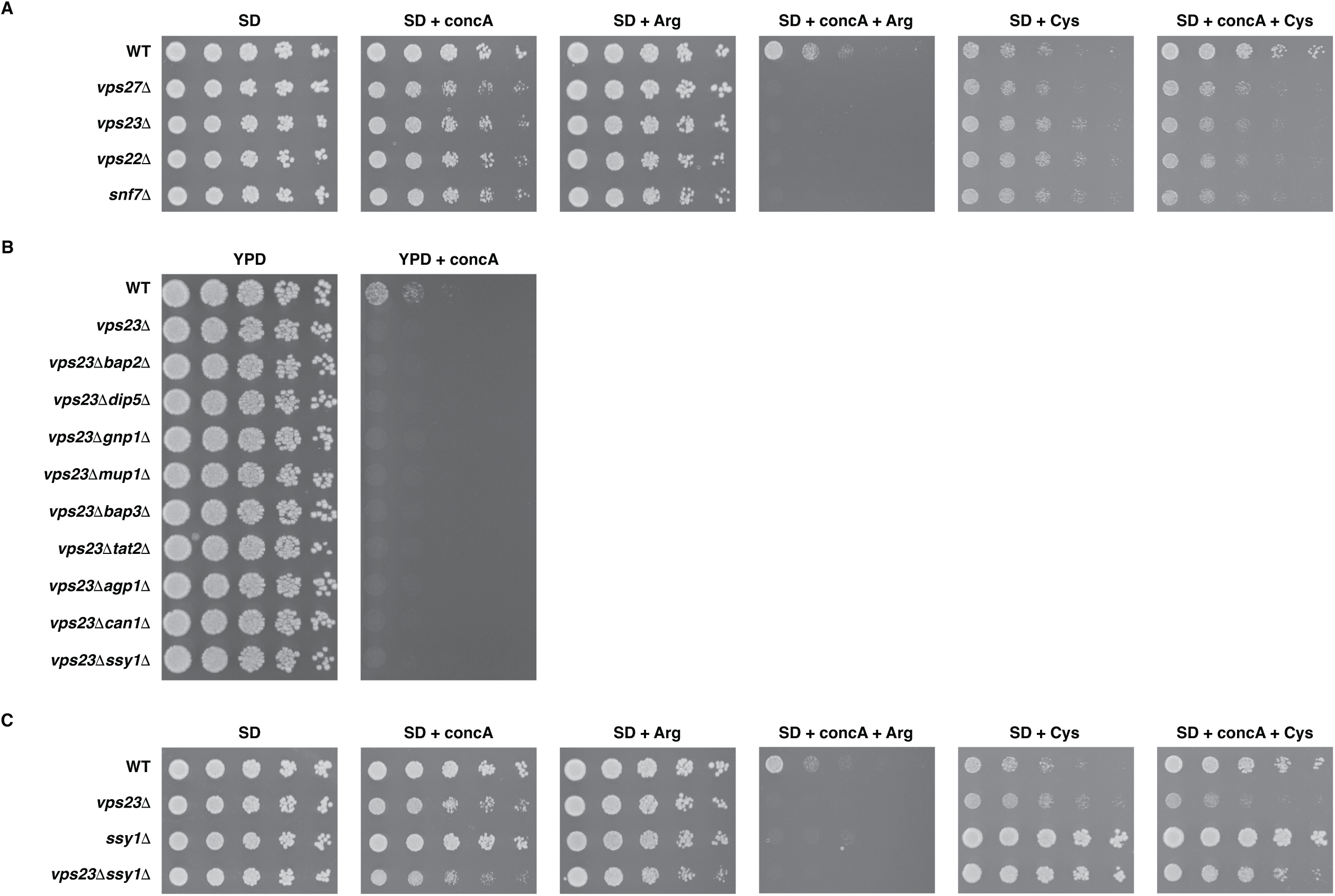
Amino acid availability drives synthetic lethality between vacuole deacidification and ESCRT disruption (related to Figure 3) (A) Spot dilution growth assays of WT and ESCRT mutant strains grown on SD medium containing DMSO or concA and supplemented with arginine (7 mM) or cysteine (7 mM). (B) Spot dilution growth assays of WT, *vps23Δ*, single plasma membrane amino acid transporter deletion strains in the *vps23Δ* background, and *vps23Δ ssy1Δ* strains grown on YPD containing DMSO or concA. (C) Spot dilution growth assays of WT, *vps23Δ*, *ssy1Δ*, and *vps23Δ ssy1Δ* strains grown on SD medium containing DMSO or concA and supplemented with arginine (7 mM) or cysteine (7 mM).

**Supplemental Figure 3.**
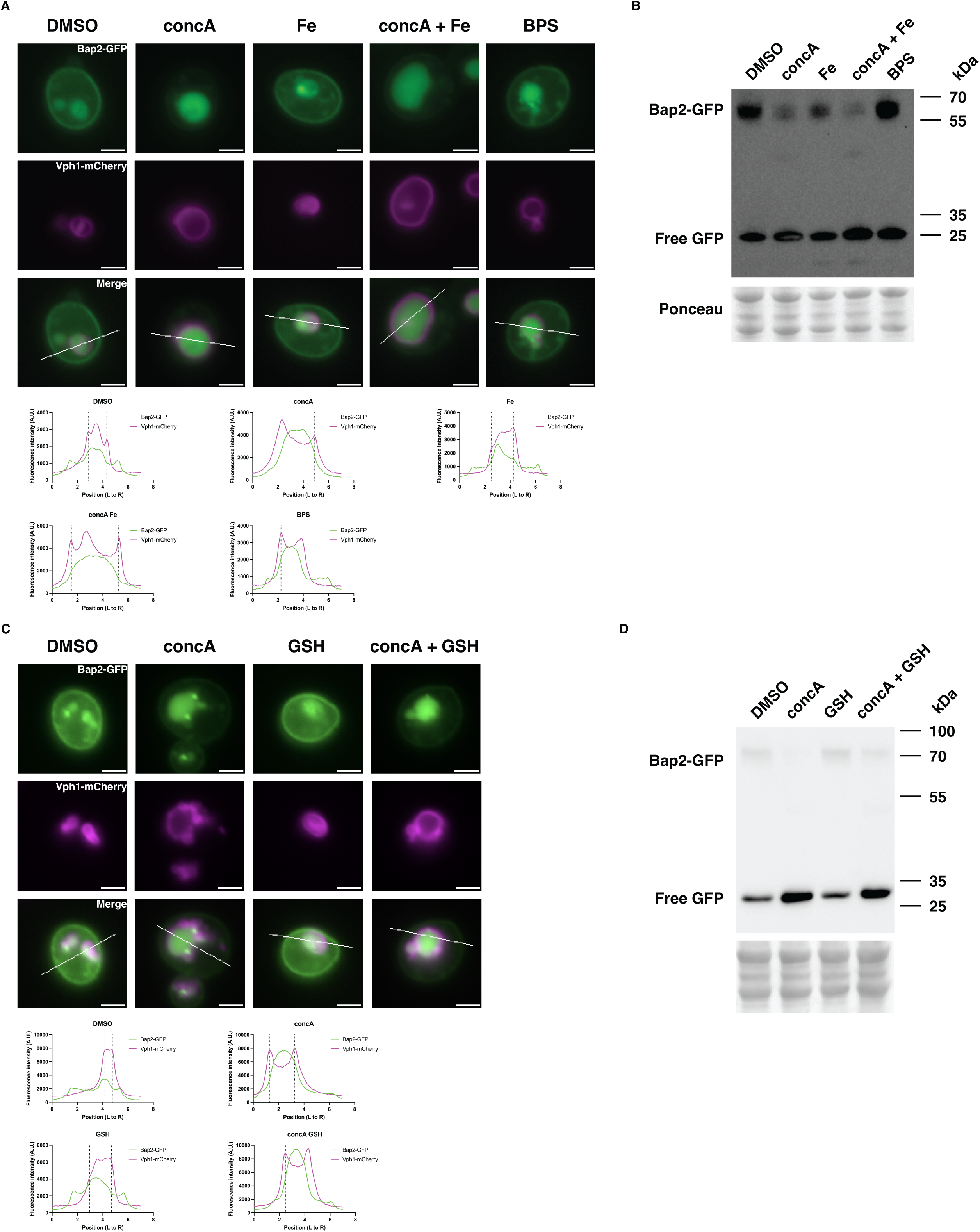
Iron limitation and oxidative stress do not drive transporter rerouting during vacuole stress (related to Figure 4) (A) Microscopy of Bap2–GFP following 3 h treatment with DMSO, concA, iron supplementation, or the iron chelator BPS in YPD medium. Scale bars: 2 µm. (B) Western blot analysis corresponding to (A). Primary antibody: α-GFP. (C) Microscopy of Bap2–GFP following 3 h treatment with DMSO, concA, iron supplementation, or the antioxidant glutathione (GSH) in YPD medium. Scale bars: 2 µm. (D) Western blot analysis corresponding to (C). Primary antibody: α-GFP.

**Supplemental Table 1. List of genes identified as hits from concA screen**

**Supplemental Table 2. Strains used in this study**

**Supplemental Table 3. Oligos used in this study**

**Supplemental Table 4. Plasmids and reagents used in this study**

